# Decorating Microbially Produced Protein Nanowires with Peptide Ligands

**DOI:** 10.1101/590224

**Authors:** Toshiyuki Ueki, David J.F. Walker, Pier-Luc Tremblay, Kelly P. Nevin, Joy E. Ward, Trevor L. Woodard, Stephen S. Nonnenmann, Derek R. Lovley

## Abstract

The potential applications of electrically conductive protein nanowires (e-PNs) harvested from *Geobacter sulfurreducens* might be greatly expanded if the outer surface of the wires could be modified to confer novel sensing capabilities or to enhance binding to other materials. We developed a simple strategy for functionalizing e-PNs with surface-exposed peptide ligands. The *G. sulfurreducens* gene for the monomer that assembles into e-PNs was modified to add known peptide ligands at the carboxyl terminus of the monomer. Strains of *G. sulfurreducens* were constructed that fabricated synthetic e-PNs with a six-histidine ‘His-tag’ or both the His-tag and a nine-peptide ‘HA-tag’ exposed on the outer surface. Addition of the peptide ligands did not diminish e-PN conductivity. The abundance of HA-tag in e-PNs was controlled by placing expression of the gene for the synthetic monomer with the HA-tag under transcriptional regulation. These studies suggest broad possibilities for tailoring e-PN properties for diverse applications.

## Introduction

Microbially produced, electrically conductive protein nanowires (e-PNs) possess properties, and possibilities for functionalization, not found in other electronic nanowire materials ^1-5^. Diverse microorganisms in both the Bacteria ^6-10^ and Archaea ^11^ produce e-PNs. Functional analysis of *Geobacter* e-PNs have demonstrated that, when attached to cells, they serve as conduits for long-range electron exchange with other cells or minerals ^1,12^. The biological roles of other microbial e-PNs have yet to be experimentally verified.

The most intensively studied e-PNs are the electrically conductive pili of the microorganism *Geobacter sulfurreducens* ^6,13-18^. *G. sulfurreducens* assembles the thin (3 nm), long (10-30 µm) e-PNs from multiple copies of just one short (61 amino acids) monomer peptide ^6,16^. Even though they are comprised of protein, the e-PNs produced with *G. sulfurreducens* are remarkably robust. They retain function under conditions required for the fabrication of electronic materials, including stability in a range of organic solvents and temperatures greater than 100 °C ^1,19^. The *G. sulfurreducens* e-PNs are produced from renewable feedstocks. No harsh chemicals are required for e-PN production and there are no toxic components in the final product. Unlike silicon nanowires, e-PNs do not dissolve in water or bodily fluids, a distinct advantage for wearable and environmental electronic sensor applications, as well as implantable electronics. e-PNs have evolved for making cell-to-cell electrical connections ^1,12^, suggesting they may be more biocompatible than other nanowire materials. The dramatic change in e-PN conductivity in response to pH ^13,17^ suggests that they may be readily adapted for diverse sensor functions ^2^.

Fabrication of e-PNs with *G. sulfurreducens* offers tight, reproducible, and consistent control of nanowire structure and electronic properties, combined with the potential for broad possibilities in the design of novel e-PNs through genetic modification of the monomer peptide ^2^. For example, the conductivity of e-PNs produced with *G. sulfurreducens* have been tuned over six orders of magnitude (ca. 1 mS/cm to1 kS/cm) ^17,18^ by genetically manipulating the abundance of aromatic amino acids in the monomer peptide.

We considered that the potential to genetically modify e-PN structure might make it feasible to decorate the outer surface of e-PNs with short peptides. The design of synthetic *E. coli* curli fibers with peptides that confer novel functions has greatly expanded their potential applications ^20,21^. However, curli fiber structure differs significantly from e-PN structure and, unlike e-PNs, curli fibers are not intrinsically conductive. Curli fibers self-assemble outside the cell ^20,21^, whereas the pilin monomers of *G. sulfurreducens* e-PNs are assembled into filaments through a complex intracellular process ^22^.

Modeling of the *G. sulfurreducens* e-PN structure predicted that amino acids at the carboxyl terminal of the pilin monomer are likely to be exposed on the outer surface of the e-PNs ^16^. This suggested that peptides might be added at the carboxyl terminus without destroying the conductive properties of the e-PNs. If so, peptides could be introduced to confer new e-PN functionalities for sensing applications, to facilitate nanowire alignment, and to link e-PNs with other materials.

Therefore, we examined whether *G. sulfurreducens* would express e-PNs containing monomers in which peptide ligands were added to the carboxyl terminus; whether the peptide ligands introduced would be accessible on the outer surface of the e-PNs; and the potential impact of the added peptide ligands on e-PN conductivity. The results suggest that *G. sulfurreducens* e-PNs can be decorated with a diversity of outer surface peptide ligands to introduce new binding properties.

## Results and Discussion

### Wires decorated with a six-histidine ligand

In order to evaluate the possibility of displaying peptide ligands on the outer surface of e-PNs, the wild-type *G. sulfurreducens* gene for the pilin monomer (PilA) was modified (Supplementary Figure 1) to encode six histidines (i.e. a ‘His-tag’) at the carboxyl end (Figure 1a). This synthetic gene was inserted into the chromosome of *G. sulfurreducens* strain KN400 (Figure 1b), along with the gene for the protein Spc that is required for pilin monomer stability ^23^. The resultant strain, which contained genes for the wild-type PilA as well as the histidine-modified PilA pilin monomer (PilA-6His), was designated strain PilA-WT/PilA-6His. Western analysis with anti-6His antibody of cell lysates of strain PilA-WT/PilA-6His separated with SDS-PAGE revealed a single protein band at the molecular weight expected for the PilA-6His monomer (Figure 1c). There was no corresponding band in lysates of wild-type cells. Western analysis with antibody that detected wild-type PilA detected a single band in wild-type cell lysates and two bands in lysates of strain PilA-WT/PilA-6His (Figure 1c). The additional band in the strain PilA-WT/PilA-6His lysate was positioned at the higher molecular weight position detected with the anti-6His antibody.

**Figure 1.**
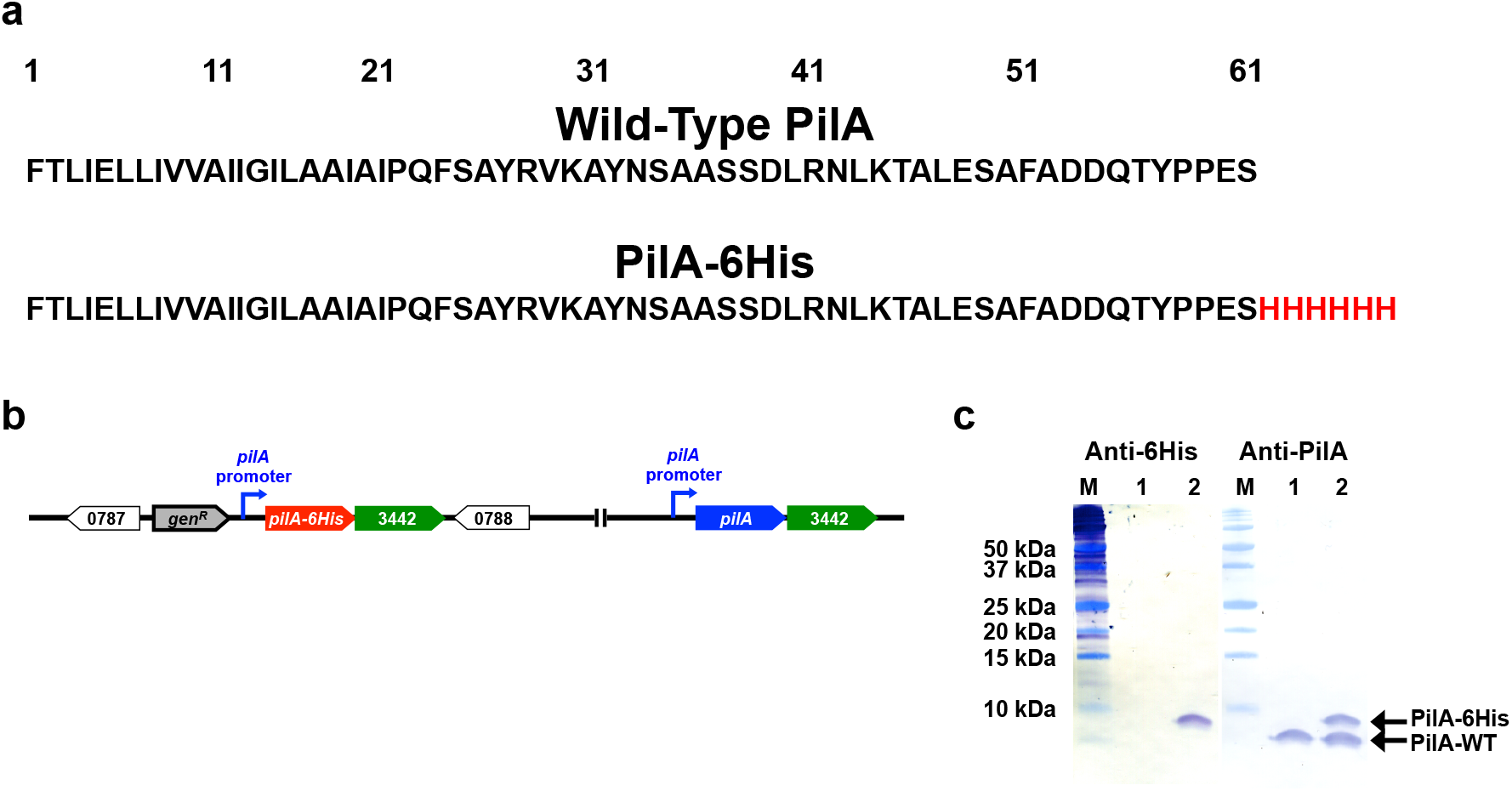
Construction of *G. sulfurreducens* strain PilA-WT/PilA-6His and expression of pilin monomers. (a) Amino acid sequence of the wild-type PilA and PilA-6His with the added histidines highlighted in red. (b) Location of wild-type and *pilA-6His* genes on the chromosome. Numbers are gene numbers from the genome sequence (NCBI, NC_017454.1). *gen*^*R*^ is the gentamicin resistance gene. (c) Western blot analysis for the wild-type (lane 1) and PilA-WT/PilA-6His (lane 2) strains. Lane M is molecular weight standard markers. Headings designate the antibody employed.

Transmission electron microscopy of cells labeled with the anti-6His antibody and a secondary antibody conjugated with gold revealed abundant His-tag loci along the wires that were accessible to the antibody (Figure 2 a, b). There was no gold labeling of wild-type cells (Supplemental Figure 2).

**Figure 2.**
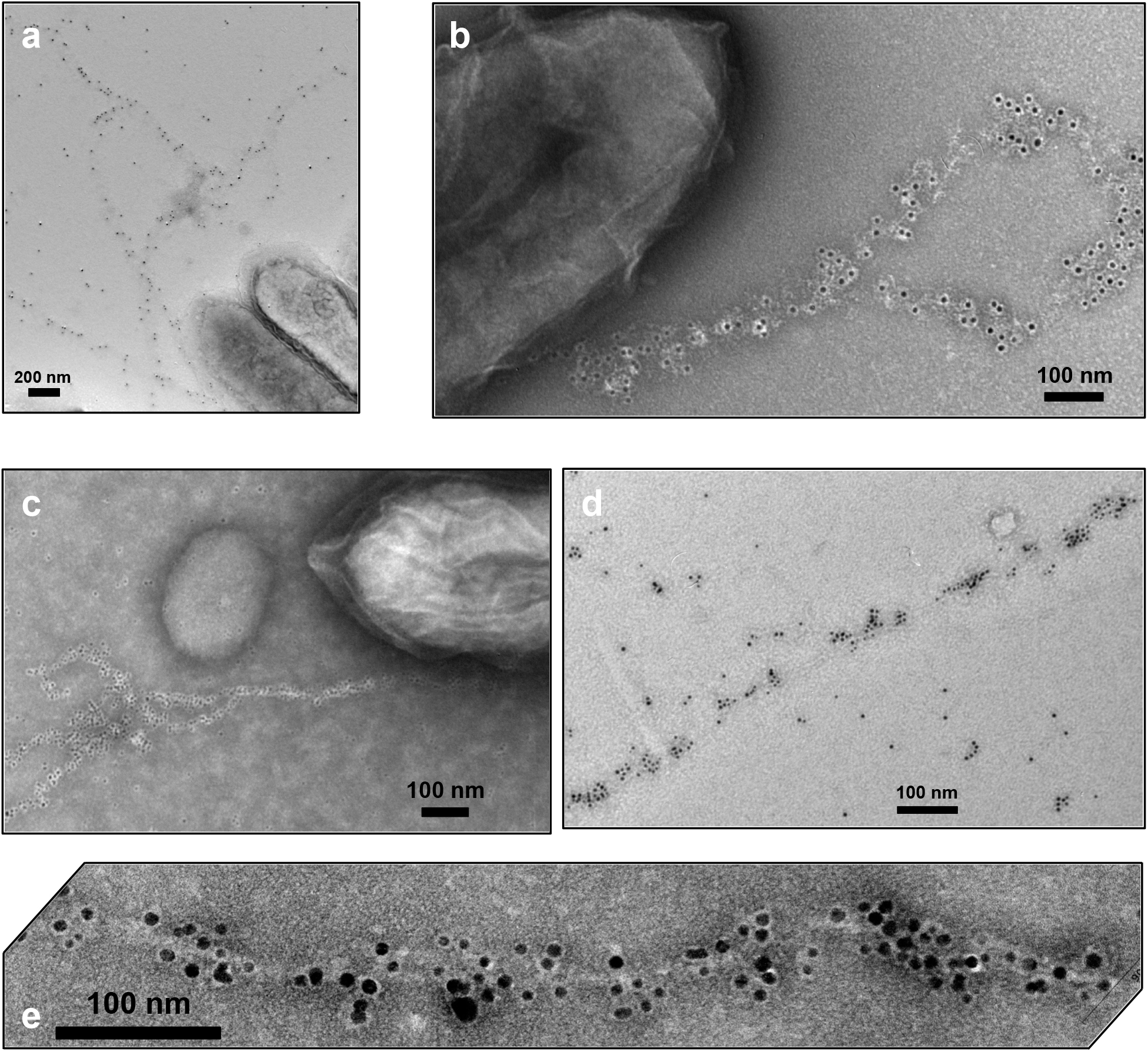
Transmission electron micrographs of immunogold (a, b) and Ni^2+^-NTA-gold (c, d, e) labeling of the 6His-tag incorporated into e-PNs of *G. sulfurreducens* strain PilA-WT/PilA-6His.

When strain PilA-WT/PilA-6His cells were treated with a Ni^2+^-NTA-gold reagent designed to label His-tags the gold nanoparticles were specifically localized along the wires (Figure 2 c,d,e). Wild-type cells were not labeled (Supplemental Figure 3). These results demonstrate that the His-tag ligand was accessible on the outer surface of the wires.

**Figure 3.**
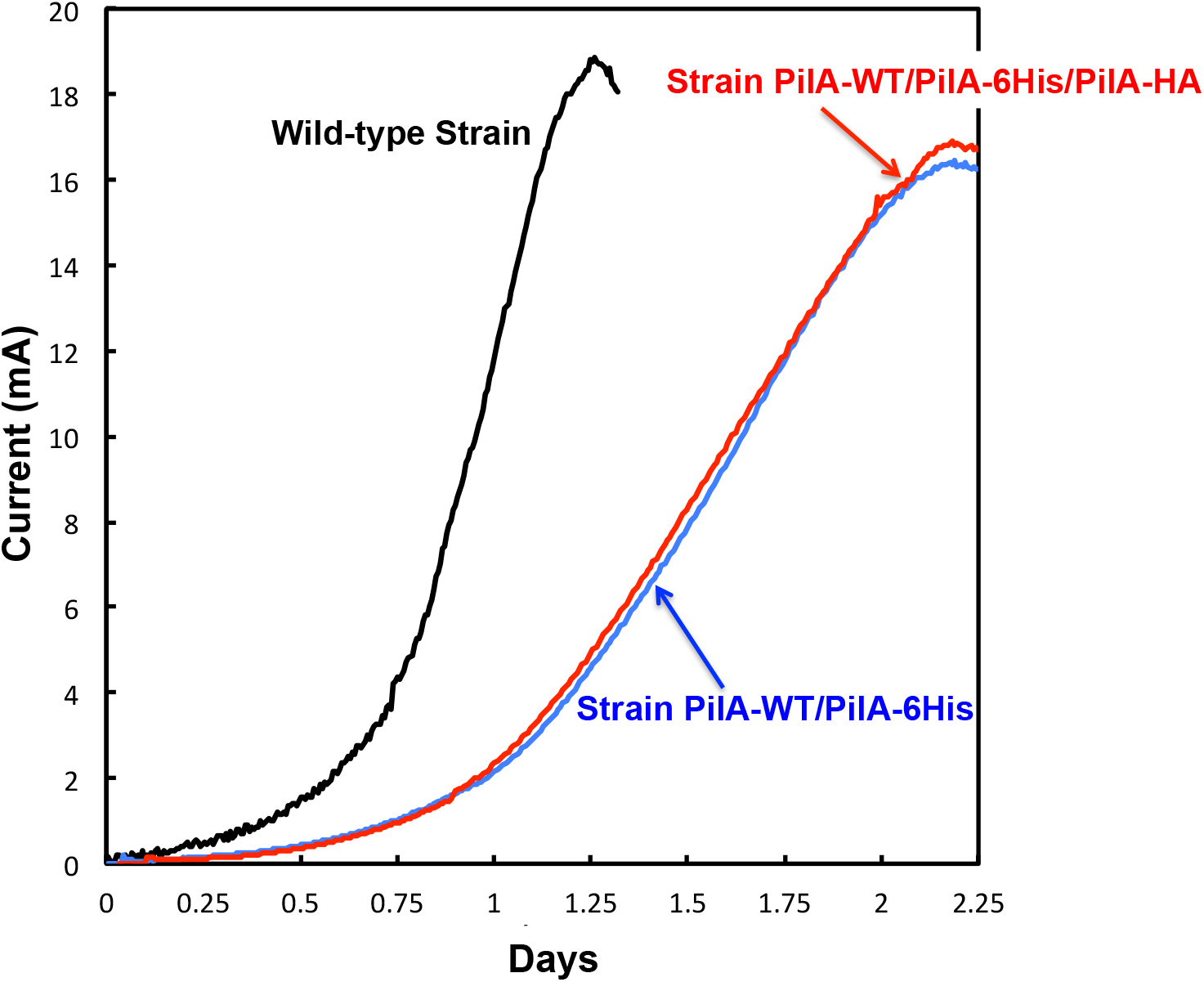
Current production of wild-type *G. sulfurreducens, G. sulfurreducens* strains PilA-WT/PilA-6His and *G. sulfurreducens* PilA-WT/PilA-6His/PilA-HA.

*G. sulfurreducens* can only produce high current densities on graphite electrodes if its pili are electrically conductive ^14,24,25^. *G. sulfurreducens* strain PilA-WT/PilA-6His produced maximum currents comparable to the wild-type strain with just a slightly longer lag in the initiation of current production (Figure 3). This result suggested that introducing the His-tag did not substantially decrease pili conductivity.

Conductivity of individual wires was more directly evaluated with conductive atomic force microscopy (c-AFM), as previously described ^11^. The wires were readily identified in topographic imaging in contact mode and had a diameter of 3.1 ± 0.3 nm (mean ± standard deviation; n= 18, 6 points on 3 wires). The conductive tip was translated to the top of the wire, and the point-mode current-voltage response (I-V) spectroscopy revealed a conductance of 7.2 + 1.5 nS (mean + standard deviation; n =9) under a load force setpoint (1 nN) (Figure 4, Supplemental Figure 4). This is slightly higher than the previously observed ^11^ conductance of 4.5 + 0.3 nS for e-PNs comprised solely of the wild-type monomer and much higher than the previously reported ^11^ conductance of the e-PNs from strain *G. sulfurreducens* strain Aro-5, which lacks key aromatic amino acids required for high conductivity.

**Figure 4.**
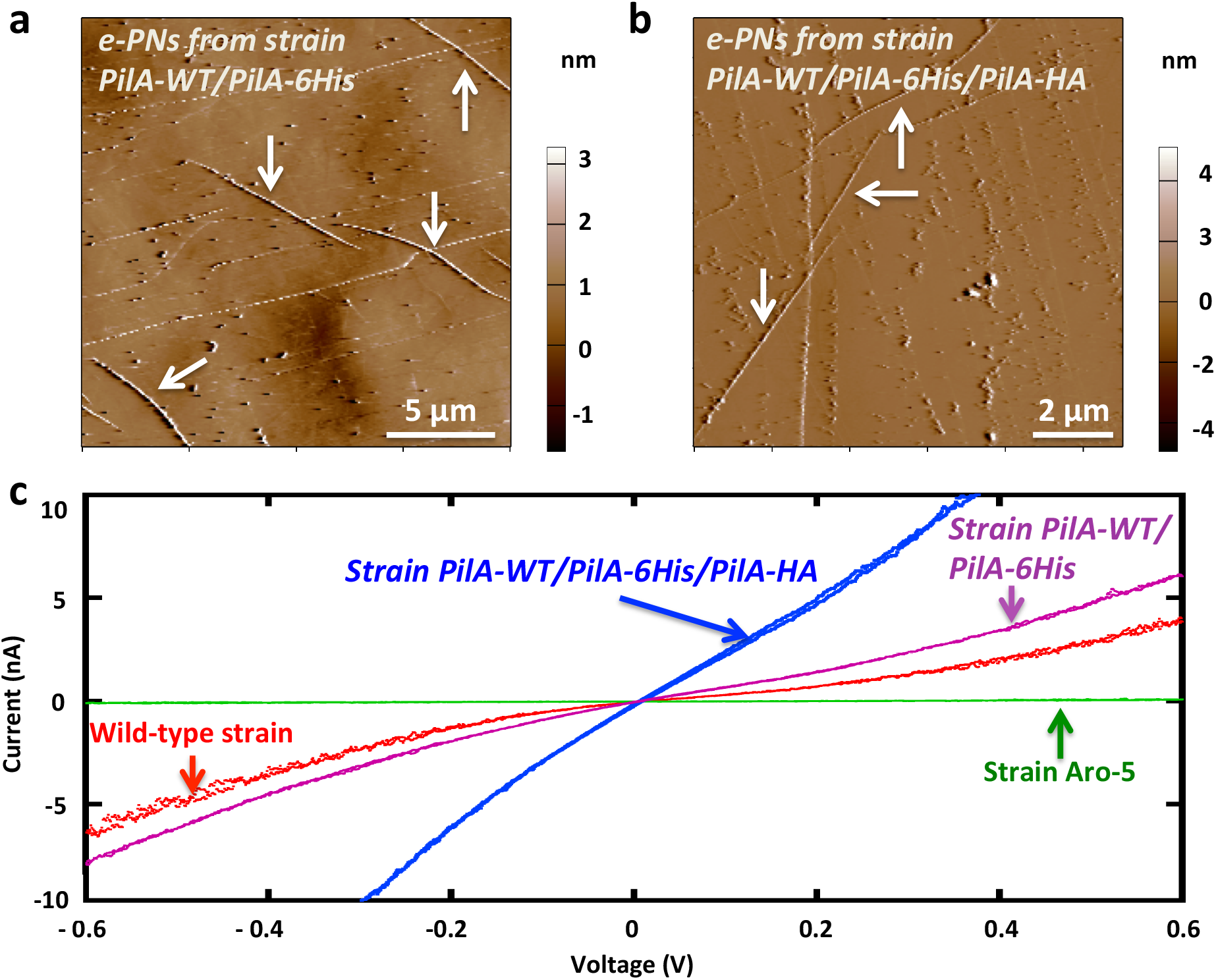
Current-voltage response of synthetic e-PNs decorated with peptide ligands. Representative contact-mode topographic imaging of e-PNs from (**a**) *G. sulfurreducens* strain PilA-WT/PilA-6His and (**b**) strain PilA-WT/PilA-6His/PilA-HA. (**c**) Point-mode current response (I-V) spectroscopy of e-PNs from PilA-WT/PilA-6His and PilA-WT/PilA-6His/PilA-HA, compared to previously published data ^11^ on e-PNs from the wild-type strain of *G. sulfurreducens* and strain Aro-5, which expresses poorly conductive pili.

### Wires decorated with two different peptide ligands

In order to determine whether two peptide ligands with different functions could be displayed on one e-PN, a gene (Supplemental Figure 1) encoding the previously described ^26^ nine-peptide ‘HA-tag’ (YPYDVPDYA) at the carboxyl end of the wild-type PilA pilin monomer (Figure 5a) was incorporated into the chromosome along with the PilA-6His and wild-type (WT) genes (Figure 5b). The gene for the PilA with the HA-tag (PilA-HA) was located downstream of the IPTG-inducible *lac* promoter/operator in order to provide the option of controlling the stoichiometry of incorporation of the PilA-HA monomer in the e-PNs (Figure 5b). This strain was designated *G. sulfurreducens* strain PilA-WT/PilA-6His/PilA-HA. Western blot analysis demonstrated that, in the presence of 1 mM IPTG, monomers of WT-PilA, PilA-6His, and PilA-HA were expressed in this strain (Figure 5c).

**Figure 5.**
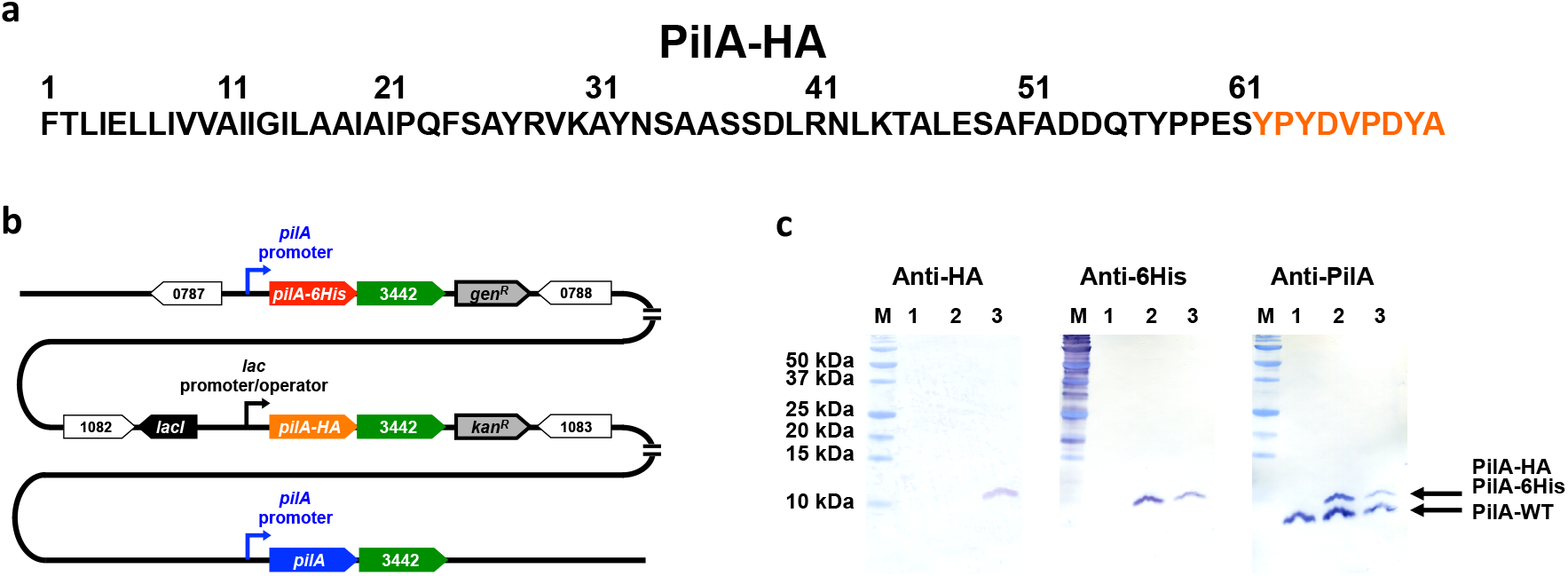
Construction of *G. sulfurreducens* strain PilA-WT/PilA-6His/PilA-HA and expression of pilin monomers. (a) Amino acid sequence of the PilA-HA with the HA tag highlighted in orange. (b) Location of wild-type, *pilA-6His*, and *pilA-HA* genes on the chromosome. *lacI* is the Lac repressor gene. *kan*^*R*^ is the kanamycin resistance gene. (e) Western blot analysis of cells lysates of the wild-type (lane 1), PilA-WT/PilA-6His (lane 2), and PilA-WT/PilA-6His/PilA-HA (lane 3) strains. Lane M is molecular weight standard markers. Headings designate the antibody employed.

Immunogold labeling for just the His-tag (Figure 6a,b) or the HA-tag (Figure 6c,d) demonstrated that both tags were abundant in the e-PNs from strain PilA-WT/PilA-6His/PilA-HA grown in the presence of 1 mM IPTG. Dual labeling with secondary antibodies with different size gold particles demonstrated that both tags were present in the same e-PNs (Figure 6 e,f). e-PNs of strain PilA-WT/PilA-6His cells were not immunogold labeled with the anti-HA antibody (Supplementary Figure 5). The current production of strain PilA-WT/PilA-6His/PilA-HA was similar to that of strain PilA-WT/PilA-6His, indicating the addition of the peptide tag did not significantly diminish pili conductivity (Figure 3). In fact, analysis of individual e-PNs of strain PilA-WT/PilA-6His/PilA-HA (Figure 4, Supplemental Figure 6) yielded higher currents at equivalent applied voltages than observed with the e-PNs with just the His-tag, with an estimated conductance of 27.2 ± 1.0 nS (n=9). A potential explanation for the higher conductance is that the HA-tag contains multiple aromatic amino acids, which may promote electron transport.

**Figure 6.**
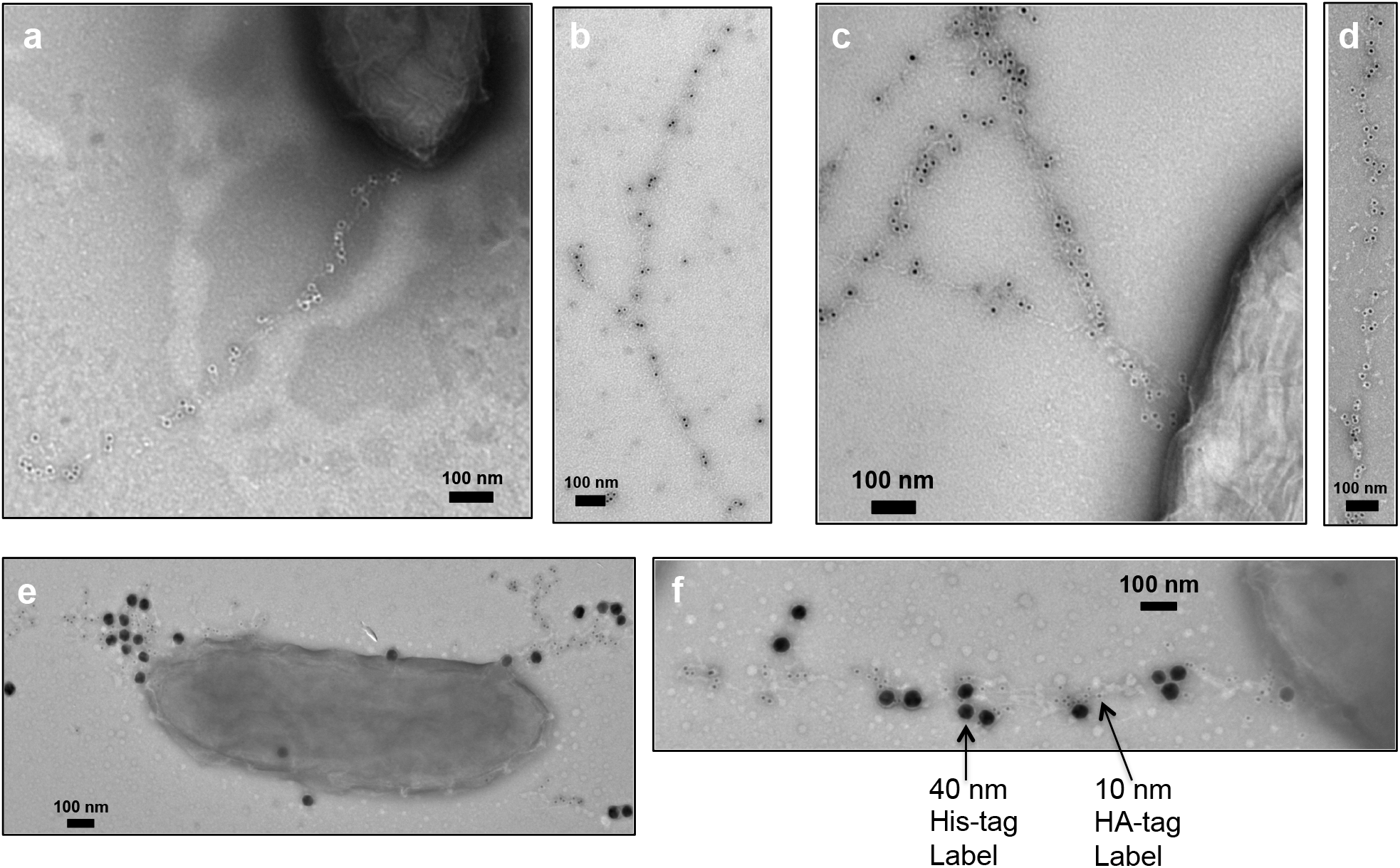
Transmission electron micrographs of immunogold labeling *G. sulfurreducens* strain PilA-WT/PilA-6His/PilA-HA for 6His-tag (a, b), HA-tag (c, d), and both tags (e, f). Gold nanoparticles were 10 nm diameter throughout with the exception of the dual labeling in panels e and f in which the 6His-tag was labeled with 40 nm diameter gold.

Some PilA-HA was expressed in strain PilA-WT/PilA-6His/PilA-HA even in the absence of the IPTG inducer (Figure 7a). However, the concentration of PilA-HA monomer in the cells was greater with added IPTG (Figure 7a). Increased pools of PilA-HA were associated with e-PNs that labeled more heavily with immunogold labeling for the HA-tag (Figure 7b,c,d). These results demonstrate that it is possible to control the abundance of a specific peptide ligand displayed on e-PNs with transcriptional control of the expression of the monomer with that ligand.

**Figure 7.**
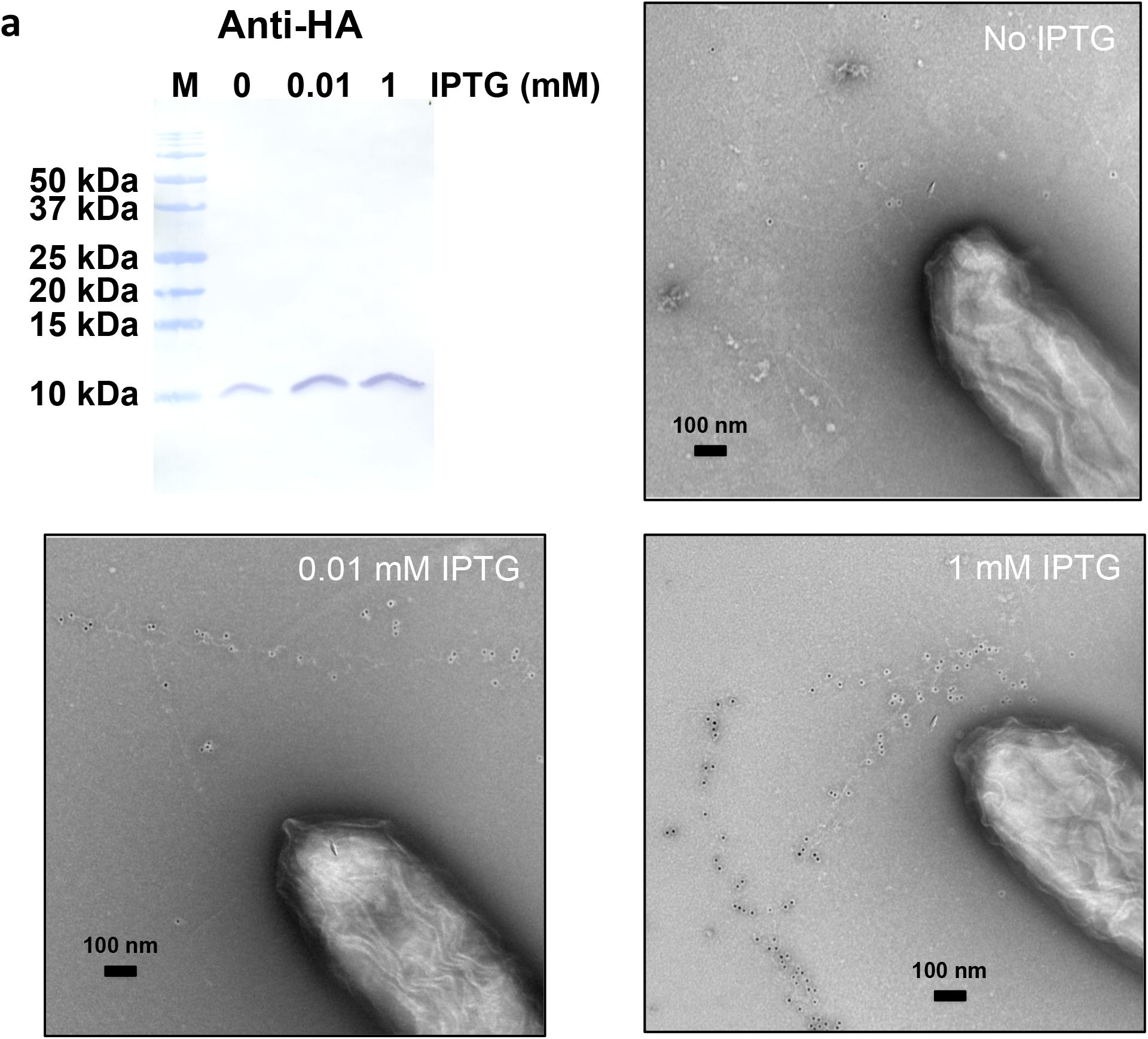
Transcriptional control of PilA-HA expression and incorporation into e-PNs in strain PilA-WT/PilA-6His/PilA-HA evaluated with anti-HA antibody. (a) Western blot analysis of cell lysates. Lane M is standard markers. (b, c, d) Immunogold labeling of HA-tag in e-PNs during growth in the presence of the designated concentration of IPTG.

### Concluding Comments

These results demonstrate that e-PNs produced with *G. sulfurreducens* can be decorated with one or more peptide ligands while maintaining, or possibly increasing, their conductivity. The stoichiometry of ligand density can be controlled with transcriptional regulation. These capabilities greatly expand the potential applications of e-PNs in electronic devices and for the fabrication of electrically conductive composite materials.

For example, sensors developed from other nanowire materials show promise for providing highly sensitive and specific, real-time electrical response for detection of diverse chemicals and biologics ^27-29^. Analytes of interest are detected as a change in nanowire conductivity that results from changes in pH associated with the activity of enzymes incorporated into the sensors, or binding of analytes to nanowires functionalized with antibodies, peptides, or other ligands. The conductivity of *G. sulfurreducens* e-PNs has already been shown to be highly responsive to pH ^13,17^. Short peptides for binding enzymes and antibodies ^30-35^ displayed on the outer-surface of e-PNs could be an effective method for functionalizing e-PN-based sensors. Furthermore, peptides can be designed to function as ligands for a wide diversity of chemical and biological analytes or to enhance attachment to cells ^36-38^. Thus, the simplicity of modifying the peptides displayed on e-PNs, and controlling the abundance of peptide display, could provide unprecedented flexibility in nanowire sensor design not readily achieved with other nanowire materials. In a similar manner, modifying the surface chemistry of e-PNs with short peptides or unnatural amino acids may enable chemical linkages with polymers or enhance binding to materials to aid in e-PN alignment in electronic devices ^1^.

The studies described here demonstrate that peptides of up to 9 amino acids can be added to the 61 amino acid monomer backbone of *G. sulfurreducens* e-PNs. However, it may be possible to decorate e-PNs with much larger peptides because the monomers of other conductive pili have an N-terminal end homologous to the *G. sulfurreducens* monomer, but are comprised of over 100 amino acids ^10,39^. These broad possibilities for modifying e-PNs with peptides coupled with the advantages of e-PNs as a ‘green’ sustainable material suggest that further investigation into the development of e-PN-based electronic devices and materials is warranted.

## Materials and Methods

### Strains and growth conditions

*G. sulfurreducens* strains were grown under anaerobic conditions at 30°C in a defined medium with acetate as the electron donor and fumarate as the electron acceptor as previously described ^40^, unless otherwise described. *Escherichia coli* was cultivated with Luria-Bertani medium with or without antibiotics ^41^.

### Construction of ***G. sulfurreducens* PilA-WT/PilA-6His strain**

*G. sulfurreducens* PilA-WT/PilA-6His strain was constructed with *G. sulfurreducens* KN400 ^42^. A gene for PilA-6His, the gene KN400_3442, which is located downstream of the *pilA* gene (KN400_1523) on the chromosome, and the putative transcription terminator were integrated at a non-coding region between KN400_0788 and KN400_0787 in the chromosome (Figure 1b). The primer pair upKpnI (CTAGGTACCGTGGTGGACCCCCTTACCGGT)/upSpeI (CGAACTAGTTGTGACCGCTGCCGGCTCCG) was used to amplify by PCR ca. 550 bp of KN400_0788 upstream of the integration site with the genomic DNA as template. This PCR product was digested with KpnI/SpeI and ligated with the vector pCR2.1Gm^r^*loxP* ^43^, resulting in pCR2.1UP-Gm^r^*loxP*. The 6His tag was fused to the C-terminal end of PilA by PCR with the primer pair pilAdnNotI (CACGCGGCCGCAAGAGGAGCCAGTGACGAAAATC)/pilACHis (GAGTTAGTGGTGGTGGTGGTGGTGACTTTCGGGCGGATAGGTTTGATC). For the construction of *pilA-6His*-KN400_3442, two PCR products were generated before being combined by recombinant PCR. *pilA-6His* was amplified with the primer pair pilAdnNotI/pilAHisrecup (CTCCAGTATGTATTTAATCAATTAGTGGTGGTGGTGGTGGTG) while KN400_3442 was amplified with the primer pair pilAHisrecdn (CACCACCACCACCACCACTAATTGATTAAATACATACTGGAG)/GSU1497XhoIAvrII (CTGCTCGAGGATACCTAGGCTATTCCGACAACTACGAGAC). The primer pair pilAdnNotI/GSU1497XhoIAvrII was then used to amplify *pilA-6His*-KN400_3442 by recombinant PCR. *pilA-6His*-KN400_3442 was cloned at NotI/XhoI sites in pCR2.1UP-Gm^r^*loxP* downstream of Gm^r^*loxP*, resulting in pCR2.1UP-Gm^r^*loxP-*pilAHis-3442. The primer pair dnAvrII (CATCCTAGGAGGGCAGACATTGCGGAACGT)/dnXhoI (CATCTCGAGCGGGTTCCGCTGCCGTCGTAC) was used to amplify by PCR ca. 530 bp of KN400_0787 downstream of the integration site. This PCR product was cloned at AvrII/XhoI sites in pCR2.1UP-Gm^r^*loxP-*pilAHis-3442, resulting in pCR2.1UP-Gm^r^*loxP-*pilAHis-3442-DN. The final plasmid was linearized with XhoI for transformation as previously described ^40^. Transformants were selected with the medium containing gentamicin (20 µg/ml) and were verified by PCR.

### Construction of ***G. sulfurreducens* PilA-WT/PilA-6His/PilA-HA strain**

*G. sulfurreducens* PilA-WT/PilA-6His/PilA-HA strain was constructed by introducing a gene encoding PilA monomer with the HA tag (*pilA-HA*) together with the gene KN400_3442 in the chromosome of the *G. sulfurreducens* PilA-WT/PilA-6His strain (Figure 5b). The *pilA-HA* gene was amplified by PCR with a primer pair TCTGGATCCAGGAGGAGACACTTATGCTTCAGAAAC/GTATTTAATCAATTACGCGTAGTCCGGCACGTCGTACGGGTAACTTTCGGGCGGATAG. The gene KN400_3442 was amplified by PCR with a primer pair CTATCCGCCCGAAAGTTACCCGTACGACGTGCCGGACTACGCGTAATTGATTAAATAC/TCTGAATTCCGATATGACTACTGCGAC. The *pilA-HA* gene and the KN400_3442 gene were connected by PCR with a primer pair TCTGGATCCAGGAGGAGACACTTATGCTTCAGAAAC/TCTGAATTCCGATATGACTACTGCGAC. The PCR product of *pilA-HA*/KN400_3442 was digested with BamHI/EcoRI and cloned in the plasmid pKIkan, which is a derivative of pKIapr ^44^ and has a kanamycin-resistance gene instead of the apramycin-resistance gene. The sequences of KN400_1082 and 1083 used for homologous recombination for introduction of *pilA-HA*/KN400_3442 are same as those of GSU1106 and 1107 for homologous recombination sequences in pKIkan, respectively. The plasmid thus constructed was linearized with XhoI for electroporation.

### Western blot analysis

The wild-type, PilA-WT/PilA-6His, and PilA-WT/PilA-6His/PilA-HA strains were grown with acetate and fumarate at 25 °C. IPTG was added at 1 mM for the PilA-WT/PilA-6His/PilA-HA strain with the exception of the study of the impact on IPTG concentrations on incorporation of PilA-HA in filaments. Cell extracts were prepared with B-PER Complete Bacterial Protein Extraction Reagent (Thermo Fisher Scientific) and the amount of protein was measured with the Bradford Protein Assay (Bio-Rad) as instructed by the manufacturer. Cell extracts were separated on 16.5% Tris-Tricine gel (Bio-Rad). An anti-PilA antibody was obtained against peptide, ESAFADDQTYPPES, corresponding to the C-terminal end of PilA (New England Peptide). An anti-6His antibody (6x-His Tag Polyclonal Antibody) and an anti-HA antibody (HA Tag Polyclonal Antibody) were purchased from Invitrogen. Western blot analysis was conducted as described previously ^45^.

### Immunogold labeling

The strains were grown with acetate and fumarate at 25°C. The PilA-WT/PilA-6His/PilA-HA strain was grown with 1 mM IPTG unless otherwise specified. Immunogold labeling was conducted as previously described ^45^. For immunogold labeling of just one type of ligand, the 6x-His Tag Polyclonal Antibody or HA Tag Polyclonal Antibody was the primary antibody and the anti-rabbit IgG-gold (10 nm) antibody (Sigma-Aldrich) was the secondary. Dual immunogold labeling was conducted with 6x-His Tag Monoclonal Antibody (Invitrogen) and the HA Tag Polyclonal Antibody as primary antibodies and an anti-mouse IgG-gold (40 nm) antibody (40 nm Goat Anti-Mouse IgG gold conjugate, Expedeon) and the anti-rabbit IgG-gold (10 nm) antibody as secondary antibodies. Samples were examined with transmission electron microscopy as described previously ^7^.

### Ni^2+^-binding assay

The wild-type and PilA-WT/PilA-6His strains were grown with acetate and fumarate at 25°C. Ni^2+^-binding assay was conducted with Ni-NTA-Nanogold (5 nm) (Nanoprobes). Seven µl of the culture was placed on a copper grid and incubated for 5 min. The grid was floated upside down in phosphate-buffered saline (PBS) for 5 min, in PBS containing 3% bovine serum albumin (BSA) and 40 mM imidazole for 15 min, and in PBS containing 0.3% BSA, 40 mM imidazole, and the Ni-NTA-Nanogold for 30 min at room temperature. The grid was washed with PBS containing 40 mM imidazole three times and with water once. Samples were stained with 2% uranyl acetate and examined by transmission electron microscopy as described previously ^7^.

### Current production

The capacity to produce current was determined in the two-chambered H-cell system with a continuous flow of medium with acetate (10 mM) as the electron donor and graphite stick anode (65 cm^2^) poised at 300 mV versus Ag/AgCl as the electron acceptor as described previously ^46^.

### Conductive atomic force microscopy (c-AFM) of individual e-PNs

As previously described ^11^, an aliquot (100 µl) of cell culture was drop-cast onto highly oriented pyrolytic graphite (HOPG). Conductive atomic force microscopy was preformed using an Oxford Instruments/Asylum Research Cypher ES atomic force microscope with a Pt/Ir-coated Arrow-ContPT tip (NanoWorld AG, Neuchâtel, Switzerland). Topographic imaging was performed in contact mode with a force of 0.1 nN. Point-mode current-voltage response (I-V) spectroscopy was achieved by applying a 1 nN force to the top of the wire and conducting quadruplicate voltage sweeps of −0.6 to 0.6 V at 0.99 Hz. The voltage sweep was averaged for each of the I-V curves and conductance was calculated from the linear portion of the I-V curve (−0.2 to 0.2 V) (Supplemental Figure 4). Average conductance and standard deviation were calculated using 3 independent points on 3 independent e-PNs of each strain. Average height and standard deviation were calculated from 6 independent points on 3 independent e-PNs.

