## Supplementary Figures for "Decorating Microbially Produced Protein Nanowires with Peptide Ligands"

**Wild-type *pilA***
ATGCTTCAGAAACTCAGAAACAGGAAAGGTTTCACCCTTATCGAGCTGCTGATCGTCGTT
GCGATCATCGGTATTCTCGCTGCAATTGCGATTCCGCAGTTCTCGGCGTATCGTGTCAAG
GCGTACAACAGCGCGGCGTCAAGCGACTTGAGAAACCTGAAGACTGCTCTTGAGTCCGCA
TTTGCTGATGATCAAACCTATCCGCCCGAAAGTTAA

***pilA-6His***
ATGCTTCAGAAACTCAGAAACAGGAAAGGTTTCACCCTTATCGAGCTGCTGATCGTCGTT
GCGATCATCGGTATTCTCGCTGCAATTGCGATTCCGCAGTTCTCGGCGTATCGTGTCAAG
GCGTACAACAGCGCGGCGTCAAGCGACTTGAGAAACCTGAAGACTGCTCTTGAGTCCGCA
TTTGCTGATGATCAAACCTATCCGCCCGAAAGT**CACCACCACCACCACCAC**TAA

***pilA-HA***
ATGCTTCAGAAACTCAGAAACAGGAAAGGTTTCACCCTTATCGAGCTGCTGATCGTCGTT
GCGATCATCGGTATTCTCGCTGCAATTGCGATTCCGCAGTTCTCGGCGTATCGTGTCAAG
GCGTACAACAGCGCGGCGTCAAGCGACTTGAGAAACCTGAAGACTGCTCTTGAGTCCGCA
TTTGCTGATGATCAAACCTATCCGCCCGAAAGTT**ACCCGTACGACGTGCCGGACTACGCG**TAA

**Figure S1. DNA sequence of wild-type *pilA*, *pilA-6His,* and *pilA-HA* genes.** Sequences for the His-tag and HA-tag are bolded in red and orange, respectively.

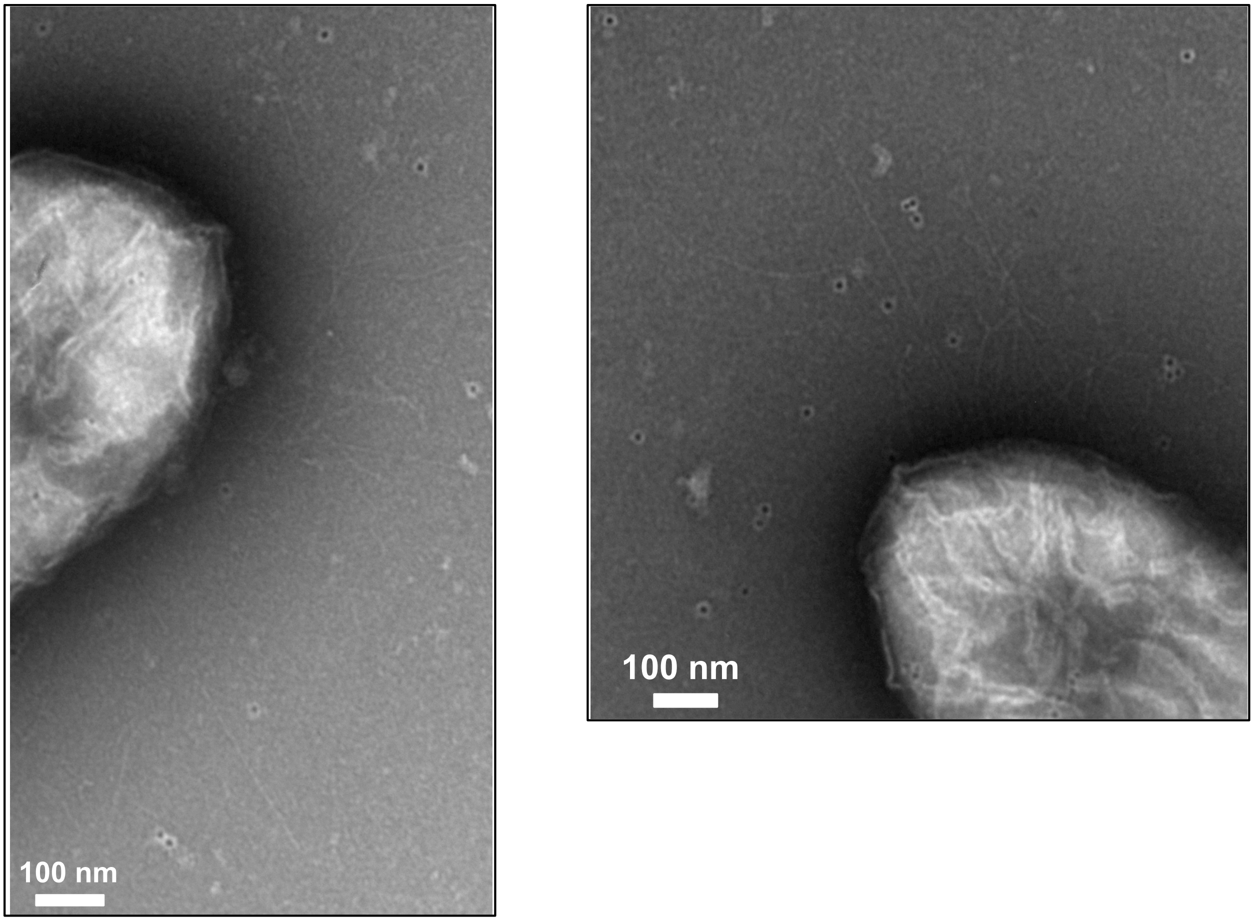


**Figure S2. Transmission electron micrographs of the wild-type strain demonstrating lack of filaments labeling with procedure for immunogold labeling of the 6His-tag.**

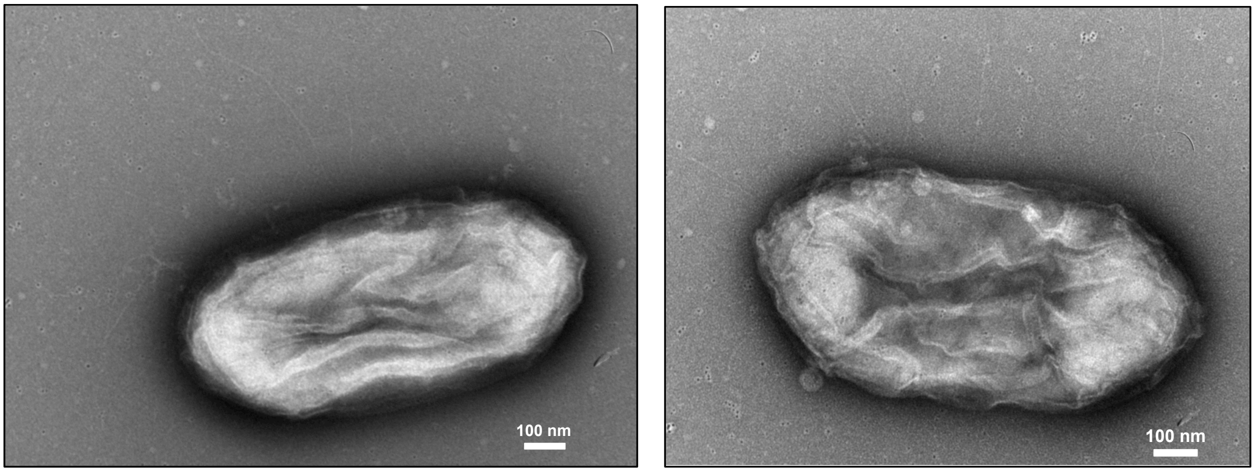


**Figure S3. Transmission electron micrographs of the wild-type strain demonstrating lack of filament labeling with the procedure for labeling the 6His-tag with $Ni^{2+}$-NTA-gold.**

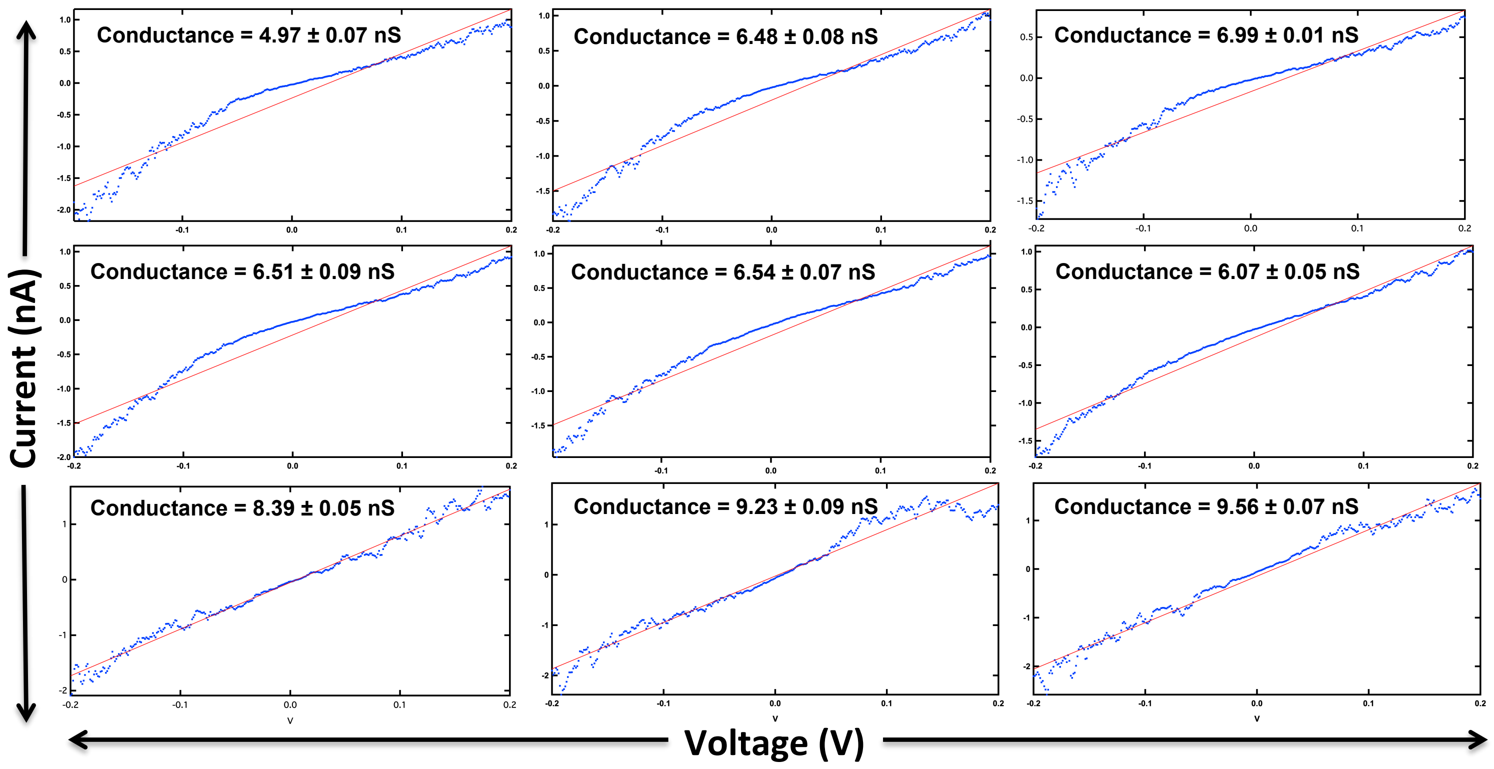


**Figure S4. Conductance for *G. sulfurreducens* strain PilA-WT/PilA-6His using point-mode current response (I-V) spectroscopy.** Three individual wires (a-c) were measured at three independent points (1-3). Calculations were made using a linear fit model between -0.2 V and 0.2 V and conduction reported as nano-Siemens (nS).

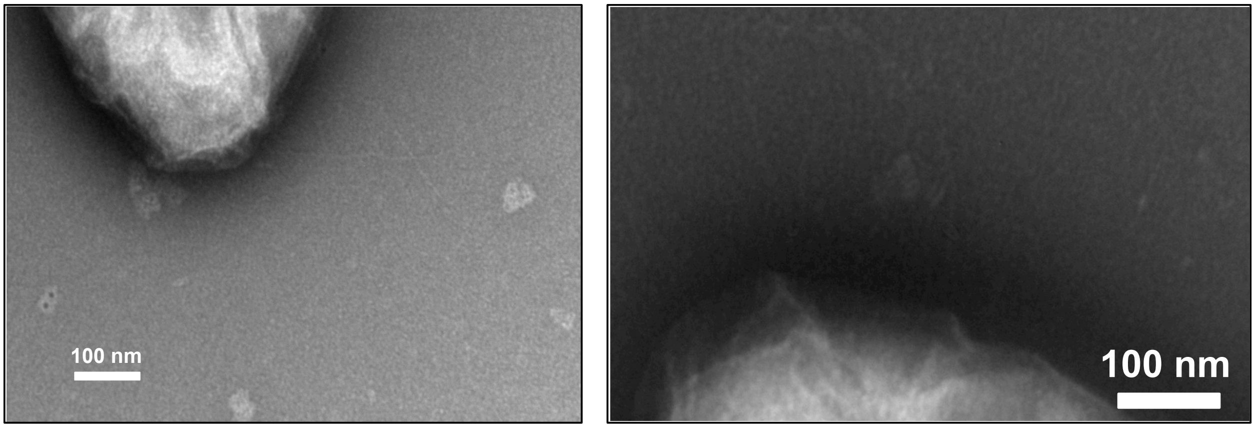


**Figure S5. Transmission electron micrographs of the PilA-WT/PilA-6His strain demonstrating the lack of labeling with the procedure for immunogold labeling of the HA-tag.**

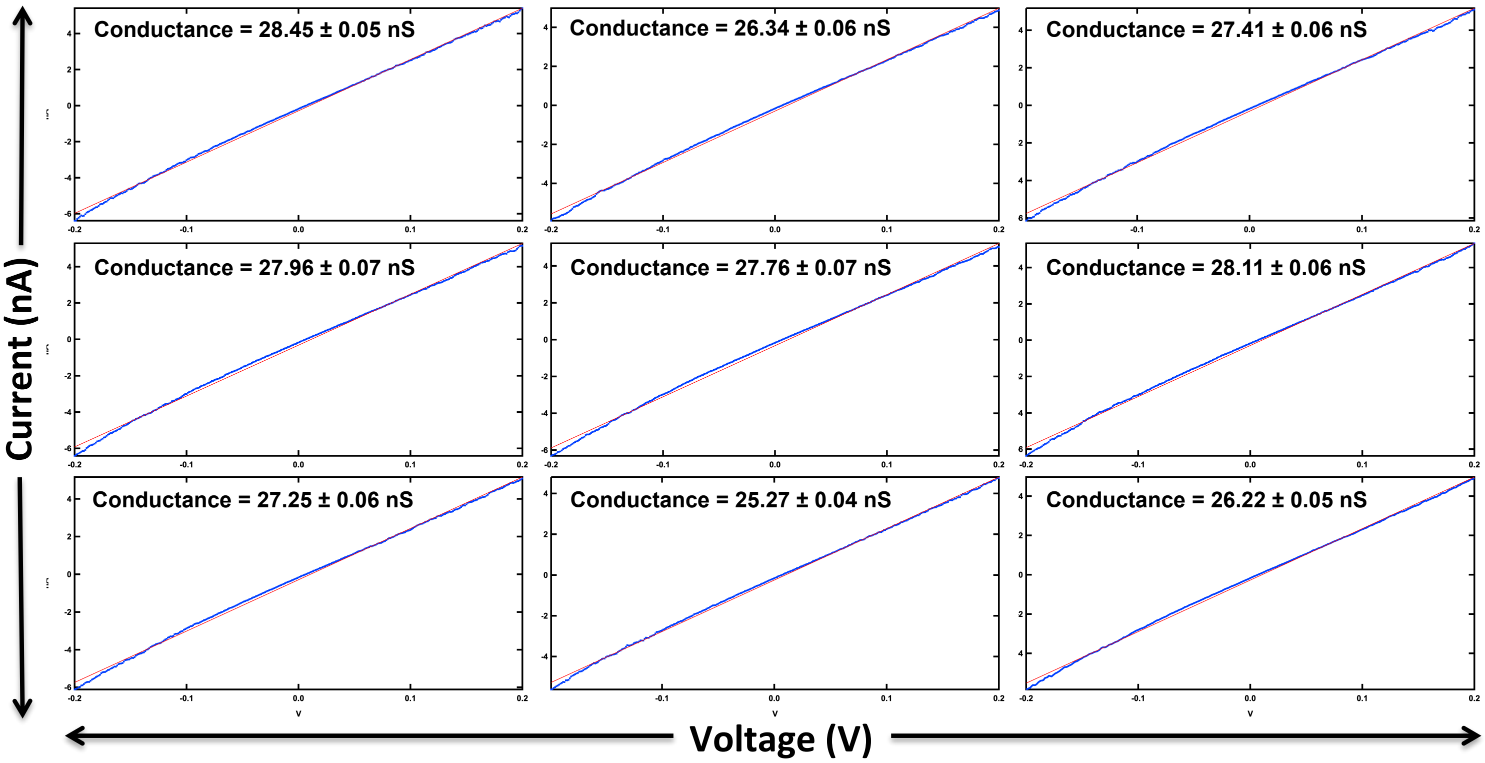


**Figure S6. Conductance for *G. sulfurreducens* strain PilA-WT/PilA-6His/PilA-HA filaments using point-mode current response (I-V) spectroscopy.** Three individual wires (a-c) were measured at three independent points (1-3). Calculations were made using a linear fit model between -0.2 V and 0.2 V and conduction reported as nano-Siemens (nS).
